# Single-cell-resolved frequency and modes of phage interactions with prokaryoplankton in the tropical surface ocean

**DOI:** 10.64898/2026.08.25.747067

**Authors:** Julia M. Brown, Alaina R. Weinheimer, Nicole Poulton, Ramunas Stepanauskas

**Affiliations:** Bigelow Laboratory for Ocean Sciences, East Boothbay, Maine; Aalborg University, Aalborg, Denmark

**Keywords:** . Phage, marine viruses, microbiology, virology, viral ecology, virus-host interactions

## Abstract

Marine planktonic viruses play critical roles in shaping microbial communities and driving global biogeochemical cycles. However, quantitative, microbiome-wide analyses of marine prokaryoplankton- virus interactions *in situ* and their ecological impacts remain challenging due to the vast diversity of viral genomes and interaction modes and the limitations of existing methodologies. Here, we utilized GORG-Tropics, a global collection of 12,715 single amplified genomes (SAGs) generated from randomly sampled marine prokaryoplankton cells, to determine the frequency and modes of their interactions with viruses in the tropical surface ocean. We found 4.2% (range 1%-19% among samples) of GORG-Tropics SAGs to contain phage genomic material, with the highest frequency found in productive ocean regions. Prokaryoplankton lineages known to have high metabolic rates, including *Prochlorococcus* and Rhodobacterales, had a substantially larger fraction of cells associated with viruses (10-12% of SAGs) as compared to the less active but highly abundant lineages including Pelagibacterales (2.2% of SAGs). Cell-virus associations indicative of lysogeny were elevated in Alphaproteobacteria relative to other taxa. The collection of phages recovered from individual SAGs exhibited genomic diversity that bridged order-level taxonomies, indicating high diversity and genomic connectivity within wild phage populations. A substantial fraction of the observed cell-virus associations disagreed with the computationally predicted host identity of the virus, indicative of non-infective interactions. The extent of genetic exchange across tailed bacteriophages infecting different hosts, and connected to taxonomically distant phages provided further evidence for the role of non-infective phage entry in the lateral transfer between tailed phages. This study provides large-scale quantitative evidence of viral infection rates in the collective prokaryoplankton community across the global surface tropical ocean through large-scale identification and quantification of the specific phages, hosts, and modes of interaction at the resolution of individual cells. Our results confirm prior reports on the overall frequency of prokaryoplankton infections with viruses in the oligotrophic tropical surface ocean. Our findings uncover non-infective phage-cell associations that may be contributing to the lateral transfer of viral genes and the nutrition of marine prokaryoplankton.

## Introduction

Marine planktonic viruses are the most abundant and diverse biological entities in the global ocean [1,2]. They influence the evolutionary trajectory of host organisms, the structure and function of microbial communities [3], and global biogeochemical cycles [4,5]. The majority of these planktonic viruses are ‘phages’—viruses that infect bacteria and archaea (“prokaryoplankton”) [6]. Marine phages have been extensively observed across the surface ocean [7,8], with estimates indicating that they are responsible for the turnover of 20 - 40% of microbial biomass daily [9,10].

While great strides have been made in capturing global marine phage diversity [7,8], major challenges in observing and quantifying phage interactions with marine bacteria and archaea remain [11]. For instance, in one of the largest viral sequencing efforts, the Global Ocean Virome, only ∼3% of the > 15,000 viral contigs recovered could be linked to a predicted, but unconfirmed host [8,12]. Yet, quantitative, lineage-resolved characterization of infections and other interactions between prokaryoplankton and viruses is critical in understanding the role of marine viruses and modelling their impact within ocean ecosystems.

Single-cell genomics offers a unique opportunity to examine the interactions of viruses and microbial cells, as a cell and all associated genetic material, including viral genomic DNA, are sequenced together [13–15]. Here, we examine the presence of viral contigs within globally sourced Single Amplified Genomes (SAGs) from the tropical and subtropical surface ocean using the Global Ocean Reference Genomes – Tropics (GORG-Tropics) SAG collection [16], documenting the distribution of phage genomic sequences across diverse microbial cells. GORG-Tropics consists of over 12,000 SAGs from 28 water samples from the Atlantic and Pacific Oceans. Through surveying the entire GORG-Tropics collection for phages, we quantify the frequency of phage interactions among epipelagic prokaryoplankton and characterize the distribution and nature of phage-microbe interactions across microbial taxa. Layering SAG microbial phylogeny with phage host predictions, we observe taxa-specific interaction frequencies and further characterize these interactions based on phage genomic content and co-occurrence of microbial genomic sequence data with identified phage genomes. We identify differences in genomic features related to infection strategy, infection frequency across taxonomic groups, and cases in which a phage’s predicted host differs from the taxonomy of the cell in which the phage was found. These observed non-host interactions document a potentially common mechanism for exchange of phage genomic material across phages capable of killing or entering phylogenetically distant hosts.

## Materials and Methods

### Virus detection and confidence scheme

12,715 SAGs from the GORG-Tropics genome collection were examined for the presence of phage genomic DNA sequences. Single amplified genomes were assembled as previously described, and only curated contigs from assemblies, published previously as the GORG-Tropics SAG collection, were scanned for viruses [17]. Contigs were scanned for viral DNA using VirSorter, VirSorter2, DeepVirFinder and VIBRANT [18–21] and an in-house workflow, viruSCope as outlined previously [14]. As a part of its virus-detection workflow, viruSCope considers the ratio of read recruitment of translated reads from the Pacific Ocean Virome; POV [22] to a LineP metagenome [23,24], providing information on metagenomic representation of read recruitment onto contigs from both datasets [13,14].

Prophage sequences, as identified by virus-finding tools, and untrimmed versions of complete contigs identified as potentially viral from each workflow were then extracted from the original, pre-curated assemblies in order to recover as much viral genomic information as possible. These contigs were combined into a single collection of candidate virus contigs, totaling 14794 contigs from 6038 SAGs. Candidate viral contigs were then analyzed by CheckV [25] (Figure S1). All contigs assigned a quality score (‘checkv_quality’ of either ‘Low-quality’, ‘Medium-quality’, ‘High-quality’ or ‘Complete’) by CheckV were maintained as well as contigs with a CheckV quality of ‘Not-determined’ in which less than 40% of contig coding sequences were identified as ‘host’ according to CheckV. This reduced the number of candidate contigs to 7670, coming from 3141 SAGs. At this point, contigs within this collection were identified as integrated into the host genome if determined so by either Virsorter, Virsorter2, VIBRANT workflows, or if CheckV categorized the contig as a ‘provirus’.

Candidate viral contigs were initially annotated using DRAMv [26], with annotations considered directly from DRAMv’s “annotations.tsv” output file. Gene annotations were determined from this file as non-hypothetical annotations from gene description columns, prioritized in the following order: ’viral_hit’, ’pfam_hits’,’vogdb_text’ and ’kegg_hit’. To increase the number of annotated genes across candidate viral contigs, protein sequences from all viral candidate contigs were clustered into orthologous groups using mmseqs2 (--min-seq-id 0.6 --kmer-per-seq 80) [16], and the most common annotation within each cluster of homologous proteins was assigned to all members of the cluster. Open reading frames from all viral candidates were additionally compared to MobileOG using mmseqs2 (‘mmseqs search’ evalue < 0.001) [27], and through search for phage structural genes using PHANNs [28].

Candidate viral contigs, including extracted proviruses, were then assessed through a combination of manual and automated evaluation of annotations of protein encoding genes in order to separate confirmed viruses from other genomic elements (Figure S1B). All identified virus contigs within each SAG were considered together when identifying confirmed viruses. Groups of candidate contigs from the same SAG were confirmed phage if any of the following conditions were met: (i) they contained hits to several phage structural genes, or hits to several other key viral genes such as small or large terminase subunits using annotations from DRAMv, MobileOG, PHANNs, and consensus annotations determined by clustering similar genes together or (ii) the collection of phage contigs from an individual SAG exceeded 15kbp and either (a) exhibited sparse annotations which hit predominantly phage genes, or (b) contigs contained open reading frames with best hits to known viruses, but no structural matches. While initially assumed that all phage contigs within a SAG originated from the same virus, there were several cases in which it was apparent that a SAG contained multiple viruses, as indicated by differences in the density of annotated genes and closest hitting phage reference genomes across groups of virus candidate contigs from the same cell (e.g. some contigs would contain a high density of ORFs matching viruses of cyanobacteria, while others from the same SAG had few annotated genes with best hits to viruses of proteobacteria). In such cases, contigs were assigned to the same phage genome based on shared annotation patterns.

To weed out other mobile and defense-related elements, contigs that were not apparently phage were assigned to additional categories based on their gene content. Contigs were identified as a mobile genetic element (‘mge’) if they contained an integrase, or plasmid-associated genes as identified via comparison to the mobileOG, but few to no other phage genes. Contigs were identified as a possible defense island (‘defense’) if no integrase was identified, but defense-associated genes, such as methyltransferases or those identified through comparison to the mobileOG, were found on the contig. Candidate viral sequences were determined to actually be host genomic sequence if either (i) they possessed more ORFs with KEGG annotations than viral annotations, (ii) if contigs had 3 or more metabolic genes in a row, or (iii) if >40% of ORFs had homology to Pfam open reading frames and more genes annotated with Pfam than with the viral databases used by DRAMv. The remaining contigs maintained the label of ‘candidate’ (Figure S1B). Finally, confirmed phages were kept if the cumulative phage contig length within the SAG was greater than or equal to 10kb. Subsequent analyses focused on this final collection of high confidence phages. After the final collection of phage sequences was verified, single amplified genomes with phage contigs removed were run through GTDB-Tk version 2.6.1 using default parameters [29], leading to the taxonomics classification of 11067 SAGs belonging to the bacteria or archaea.

The SAGs that compose the GORG-Tropic collection originated from cells were selected using fluorescence activated cell sorting as described previously [16]. 10523 cells within the collection were selected based on Syto9 fluorescence (i.e. nucleic acid content) and being within a prokaryoplankton size range. From the most deeply sampled site, SWC-09, 2019 additional cells were selected based on elevated RSG fluorescence and prokaryoplankton size range. Identified phages originating from SAGs in all FACS mode categories were run through all described workflows, but only the prokaryoplankton-sized SAGs selected based on Syto9 fluorescence are included for described site- and Order-level infection trends, as these may be considered randomly sampled members of the prokaryplankton community. Phenotypic information, including Syto9 fluorescence and cell size estimates, were collected and calculated during cell sorting and previously reported by Pachiadaki et al 2019.

### Host prediction

Several methods were employed for *in silico* prediction of hosts of identified phages. We first utilized a gene homology-based approach through quantification of closest hits to RefSeq’s viral database, extracted from the “annotations.tsv” table produced by DRAMv. Predicted host was assigned if >20% of phage open reading frames most closely hit phages of a shared host or if the hits for the most common phages matched the determined phylogeny of the host cell. To further identify putative hosts, contigs from identified phage genomes were run through iPHOP version 1.4.1[30]. Not all contigs within the phages were assigned host predictions, and often there were conflicting predictions within identified phage genomes across contigs. Host predictions made by iPHOP were maintained if one host was predicted across all contigs for which a prediction was made for each identified phage, or based on the host assignment with highest confidence. iPhOP host assignments were used for those phage genomes for which a DRAMv homology-based assignment was not determined. iPHoP allowed for host prediction for an additional 224 viruses. Finally, for phages for which the above host prediction methods failed, phage host was assigned based on the microbial taxonomy of the SAG the phage contigs were found in, as determined by GTDBtk.

### Gene-specific analyses

Phages recovered from GORG-Tropics were scanned for genes encoding the cyanobacterial photosystem II D1 protein (i.e. PsbA) based on consensus annotations of identified gene clusters as described above. Translated protein sequences were aligned with *Prochlorococcus, Synechococcus* and cyanophage photosystem II D1 protein sequences extracted from using clustalOmega [31] with default alignment parameters. The alignment was filtered to keep only columns with <= 1% gaps, resulting in a final alignment of 150 amino acids for phylogenetic analysis using IQTree with default parameters [32]. The phylogenetic tree was visualized in R using the ggtree, ggplot2, ggtreeExtra and ggstar R packages.

Phages recovered from GORG-Tropics were scanned for genes encoding the tail fiber proteins by searching gene clusters for genes with a consensus annotation containing the phrase ‘fiber’ or ‘spike’ but not ‘connector’, ‘shaft’ or ‘tube’, and identification of additional predicted tail fiber proteins using PHANNs predictions (tail fiber annotations with score limit > 9)[28]. To draw collector’s curves, phages were divided by their predicted host group and sorted in descending order based on the number of identified tail fibers per phage. Phages were sampled in this order, and the number of unique accumulated tail fibers per added phage was calculated.

Statistical analyses were conducted using methods within the python packages scipy.stats, sklearn.cluster, sklearn.utils, and scikit_posthocs. Comparisons of means for phenotypic characteristics by cell infection status, and comparison of phage representation within POV by predicted host were conducted using a Kruskal-Wallis tests, followed by Dunn’s post hoc tests with bonnferoni correction using “scikit” and “statsmodels” python packages. Code used to run analyses and generate figures is available at https://github.com/juliambrosman/gorg-tropics-viruses.

### Protein sharing

Protein content of phage genomes, identified by prodigal run within the DRAMv workflow, were compared via vContact2 version 0.11.3 [33] to 22,094 phage genomes downloaded from NCBI on May 11, 2023 using the INPHARED workflow [34] to determine phylogenetic relationships of phages based on shared protein clusters. The resulting vContact2 network was visualized using Cytoscape [35] using an unweighted perfuse force-directed layout. Network statistics were calculated using Networkx [36]. Orthologous proteins identified between phages were further examined using blastp [37], keeping all hits for which 80% of the smaller sequence aligned, for reciprocal comparison of amino acid sequences of open reading frames recovered from identified phage genomes. Homologous genomic regions were visualized using clinker [38].

## Results and Discussion

### Distribution of phage interactions

Here, we examined phage-microbe associations across the surface tropical ocean by identifying phage genomic sequences within 12,542 single-amplified genomes (SAGs) sourced from prokaryoplankton-sized DNA-containing particles belonging to the GORG Tropics SAG collection. GORG-Tropics is composed of SAGs from 28 seawater samples, ranging from 180 to 6,063 SAGs per site, all with genome assembly sizes >20kb [16]. SAGs originate from prokaryoplankton- sized particles, determined using light forward scatter as a proxy for cell size during fluorescence activated cell sorting. For the most deeply sequenced site, SWC-09, 2019 cells were sorted based on an activity probe (RedoxSensor Green); we will refer to this subset of SAGs as redox-active SAGs. All other SAGs included in this study from SWC-09 and all other sites were sorted based on nucleic acid content using a DNA stain (SYTO-9); we will refer to these as randomly-sampled SAGs. Overall, 524 phages were identified in 507 SAGs from GORG-Tropics, with 4.2% of randomly sampled cells containing a phage.

The fraction of randomly sampled phage-containing SAGs ranged from < 1% to 19% of SAGs sampled per site (Figure 1A). The highest fraction of phage-containing SAGs was found in samples collected north of the Amazon River plume and off the coast of northwest Africa. Both regions have previously been shown to have enhanced productivity due to riverine inputs [39,40] and upwelling [41], respectively, suggesting that higher productivity may lead to higher overall rates of phage-microbe interactions. 14 SAGs were identified as containing multiple phages, the majority of which were found (11/14 SAGs with > 1 phage) within these two regions.

**Figure 1.**
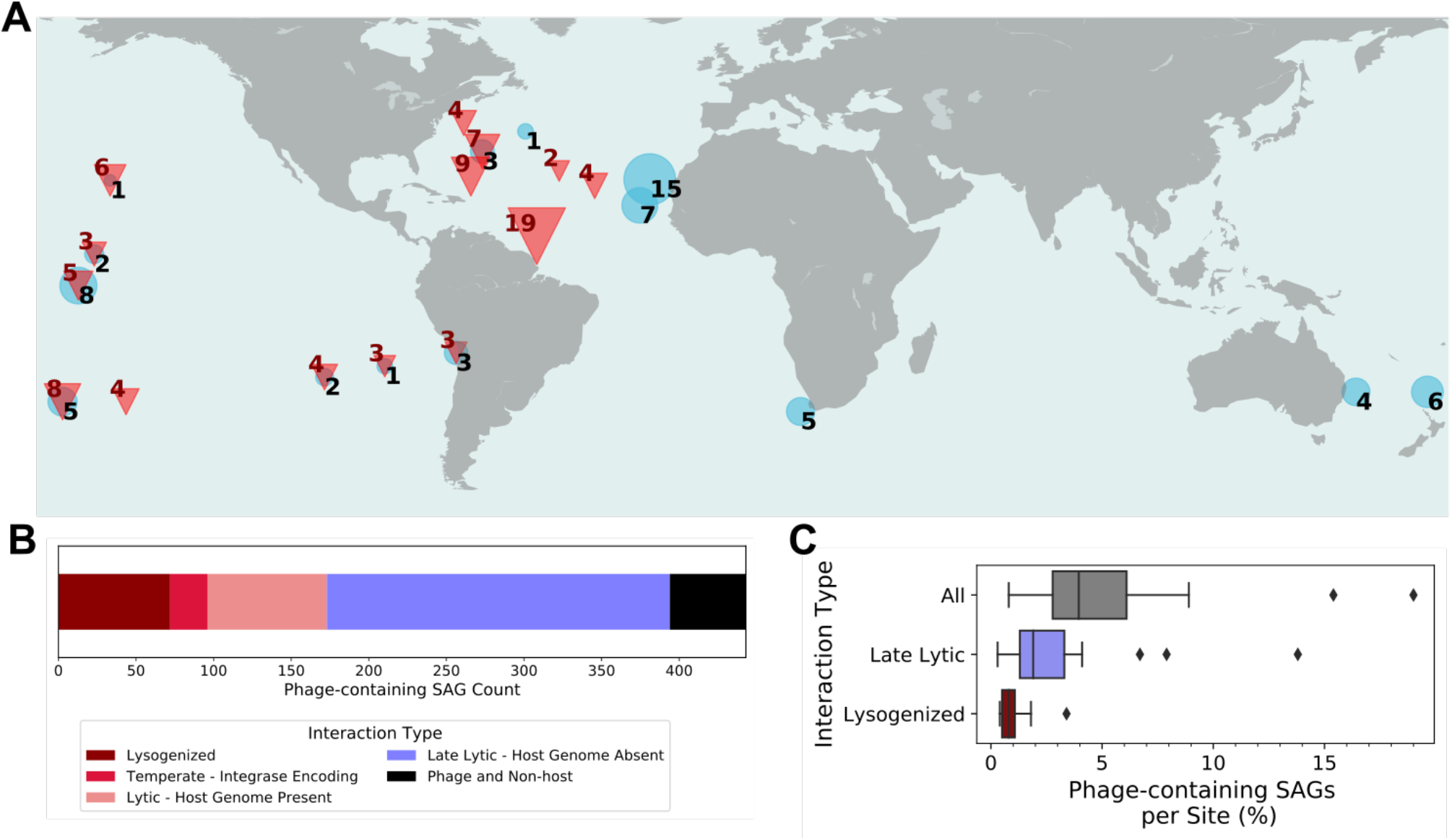
Geographic and phylogenetic distribution of viral-cell associations. A. The fraction of phage-containing randomly sampled SAGs in samples from which >50 SAGs were obtained. Data marker size and associated numerals indicate the percent of SAGs containing viral sequences in each seawater sample. Red triangles indicate the deep chlorophyll maximum or > 75m depths. Blue circles indicate depths <75m. B. Overall distribution of interaction types across randomly-sampled GORG-Tropics cells that contained a phage. C. Proportion of randomly sampled SAGs containing phage per site.

0.6% of randomly sampled cells (72 SAGs) recovered from GORG-Tropics, appeared to be lysogenized, as indicated by phage co-assembly with host genomic regions (Figure 1B), constituting 0.4 to 3.4 percent of SAGs per site (Figure 1C). While our observed prevalence of lysogeny is likely an underestimate due to the conservative approaches taken to identify high-confidence phages, this result is remarkably consistent with prior experimental reports of low to undetectable rates of lysogeny within many tropical and oligotrophic marine surface samples [42–44]. This is in contrast to rates of lysogeny observed in microbial genome collections containing non-marine organisms, such as a survey of Genbank published in 2016, which found nearly half of all available bacterial genomes to be lysogenized by a phage using similarly stringent criteria for phage identification [45]. These findings suggest that epipelagic marine prokaryoplankton contain fewer prophages than microbial groups dominating current GenBank genome deposits.

Slightly over half of randomly sampled phage-containing SAGs (234) contained no identifiable microbial genomic information, amounting to 1.8% of randomly sampled GORG-Tropics cells overall and ranging from 0.3 to 13.8% of cells per sample site (Figure 1C). We assume this characteristic is indicative of host genome degradation often seen during late stages of lytic infection, and likely coincides with the cell containing observable viral capsids. Particle optical properties measured during cell sorting indicate that these “phage only” assemblies originate from prokaryotic cells rather than free viral particles (Figure S2). We can thus compare these numbers to prior reports of the frequency of visibly infected cells (FVIC) as determined by Transmission Electron Microscopy (TEM) in a variety of marine surface samples. Our observations are consistent with prior FVIC measurements, which, for example, ranged from 0.9% to 4.3% in tropical and temperate surface samples [46]) and from undetectable to 4.2% of cells in the Adriatic Sea [47].

Phenotypic comparisons of randomly sampled cells at different stages of infection revealed differences in cell size and nucleic acid content depending on the cell’s infection state. The average estimated diameter of cells containing phage DNA was significantly larger than cells without phage present, revealing virus-induced changes in cell physiology within phage-infected cells (Figure S2A and S2C). The FACS-based cell size estimates assume a uniform diameter-to-light-scatter ratio among cells, which may bias estimates for infected cells if viral particles increase that cell’s light reflectivity. The increase in cells sizes for both late-lytic and other infection types, however, suggest an impact beyond the presence of viral particles. There were significant differences in Syto9 fluorescence between cells in different infection states, with cells in which only phage DNA was detected having the weakest Syto9 fluorescence values, and cells with both microbial and phage DNA having the highest Syto9 fluorescence values, indicating changes in total intracellular nucleic acids between infection states (Figure S2B).

### Patterns in phage-microbe interactions by host taxonomy

Given the large proportion of phage-containing cells without identifiable host genomic sequences, we combined phage-host co-occurrence observations with *in silico* host prediction techniques to determine the likely taxonomy of virus-only cells and to calculate overall frequencies of phage association across microbial taxa. Cells were assigned a microbial taxonomy based on the classification of microbial genomic sequences, if present, and alternatively by the predicted host taxonomy of identified phage contigs. We found that 11.4 % (62) of host predictions were in conflict with the taxonomy of the SAG from which the phage was recovered (Figure 1B, Table 1). We called these cases non-infecting “phage and non-host” interactions, and considered them separately from overall frequencies of order-level frequencies of phage-host interactions.

**Table 1.** Characteristics of phage associations across order-level taxa from randomly sampled GORG-Tropics SAGs that contained at least 50 SAGs within the order. Values are displayed as SAG counts per category, with percent of total cells in each order in parentheses.

| Taxon | Total in<br>GORG-Tropics | Integrated<br>Phage | Integrase-<br>encoding<br>Phage and<br>Host | Phage<br>and Host | Phage<br>Only | Non-host<br>phage<br>and<br>Microbe | Total Phage<br>Associations | Total Phage<br>Infections |
| --- | --- | --- | --- | --- | --- | --- | --- | --- |
| Pseudomonadota__Alphaproteobacteria__Pelagibacterales | 3801 | 28 (0.7%) | 7 (0.2%) | 22 (0.6%) | 18 (0.5%) | 25 (0.7%) | 100 (2.6%) | 75 (2.0%) |
| Cyanobacteriota__Cyanobacteriia__PCC-6307 | 909 | 9 (1.0%) | 5 (0.6%) | 23 (2.5%) | 80 (8.8%) | 1 (0.1%) | 118 (13.0%) | 117 (12.9%) |
| Pseudomonadota__Gammaproteobacteria__SAR86 | 897 | 0 (0.0%) | 0 (0.0%) | 0 (0.0%) | 3 (0.3%) | 1 (0.1%) | 4 (0.4%) | 3 (0.3%) |
| Bacteroidota__Bacteroidia__Flavobacteriales | 662 | 2 (0.3%) | 1 (0.2%) | 5 (0.8%) | 12 (1.8%) | 5 (0.8%) | 25 (3.8%) | 20 (3.0%) |
| Pseudomonadota__Alphaproteobacteria__HIMB59 | 571 | 2 (0.4%) | 0 (0.0%) | 1 (0.2%) | 0 (0.0%) | 3 (0.5%) | 6 (1.1%) | 3 (0.5%) |
| Actinomycetota__Acidimicrobiia__Actinomarinales | 482 | 3 (0.6%) | 1 (0.2%) | 4 (0.8%) | 5 (1.0%) | 2 (0.4%) | 15 (3.1%) | 13 (2.7%) |
| Pseudomonadota__Alphaproteobacteria__Puniceispirillales | 238 | 2 (0.8%) | 1 (0.4%) | 1 (0.4%) | 5 (2.1%) | 1 (0.4%) | 10 (4.2%) | 9 (3.8%) |
| Pseudomonadota__Alphaproteobacteria__Rhodobacterales | 203 | 10 (4.9%) | 6 (3.0%) | 4 (2.0%) | 1 (0.5%) | 0 (0.0%) | 21 (10.3%) | 21 (10.3%) |
| Pseudomonadota__Gammaproteobacteria__Pseudomonadales | 170 | 1 (0.6%) | 0 (0.0%) | 0 (0.0%) | 2 (1.2%) | 1 (0.6%) | 4 (2.4%) | 3 (1.8%) |
| Thermoplasmata__Poseidonii__Poseidoniales | 146 | 1 (0.7%) | 0 (0.0%) | 6 (4.1%) | 1 (0.7%) | 1 (0.7%) | 9 (6.2%) | 8 (5.5%) |
| Marinisomatota__Marinisomatia__Marinisomatales | 124 | 1 (0.8%) | 0 (0.0%) | 1 (0.8%) | 0 (0.0%) | 1 (0.8%) | 3 (2.4%) | 2 (1.6%) |
| Pseudomonadota__Alphaproteobacteria__TMED127 | 83 | 0 (0.0%) | 0 (0.0%) | 0 (0.0%) | 0 (0.0%) | 1 (1.2%) | 1 (1.2%) | 0 (0.0%) |
| Pseudomonadota__Gammaproteobacteria__GCA-002705445 | 55 | 0 (0.0%) | 0 (0.0%) | 0 (0.0%) | 0 (0.0%) | 1 (1.8%) | 1 (1.8%) | 0 (0.0%) |
| SAR324__SAR324__SAR324 | 51 | 4 (7.8%) | 2 (3.9%) | 0 (0.0%) | 0 (0.0%) | 0 (0.0%) | 6 (11.8%) | 6 (11.8%) |
| Pseudomonadota__Alphaproteobacteria__MED-G09 | 50 | 1 (2.0%) | 0 (0.0%) | 0 (0.0%) | 0 (0.0%) | 0 (0.0%) | 1 (2.0%) | 1 (2.0%) |
| Other | 2081 | 8 (0.4%) | 1 (0.0%) | 79 (3.8%) | 10 (0.5%) | 6 (0.3%) | 103 (4.9%) | 97 (4.7%) |

Overall frequencies of phage-host associations within randomly sampled SAGs differed between taxonomic groups (Figures 2B, S2A) with differences potentially corresponding to the activity of host cells. For example, marine *Prochlorococcus* SAGs had the highest proportion of SAGs with viral contigs (12.9%). Marine picocyanobacteria are the most transcriptionally active microbes in the oligotrophic surface ocean [48], dominate photosynthetic biomass of the open ocean and are highly efficient photosynthesizers [49]. Our observed rate of viral associations with marine cyanobacterial cells is higher than previously reported infection rates using the single cell polony method, which observed 0.35 – 1.6% of cyanobacterial cells to be infected [50]. Differences in observed infection frequencies could be due to the way in which cells are selected (random selection for GORG-Tropics versus selection based on photopigment content for ipolonies) and how phage infections were observed (untargeted genomics-based here versus PCR primer-based observations that rely on homology for detection). Members of the order Rhodobacterales also had a high frequency of phage-host associations (10.3% of cells classified as Rhodobacterales were associated with a phage). The majority of hosts belonging to the Rhodobacterales order fell within the Rhodobacteraceae family, members of the marine *Roseobacter* clade, including LGTR01 (8 SAGs) and LFER01 (6 SAGs) genera. Marine Rhodobacteraceae were previously observed to have the highest levels of respiration despite their relatively low abundances [51–54]. The higher rates of phage interactions of these two groups indicate that viruses may more frequently target metabolically active microbial community members, consistent with the kill the winner hypothesis [55,56]. Other taxa that exhibited comparably high infection frequencies include SAR324, with 6 out of 51 cells infected (11.8%), and archaeal order *Poseidoniales* (5.5% infected, Table 1), with members of the latter belonging to the genera MGIIb-P, UBA36, Thalassarchaeum and Poseidonia.

**Figure 2.**
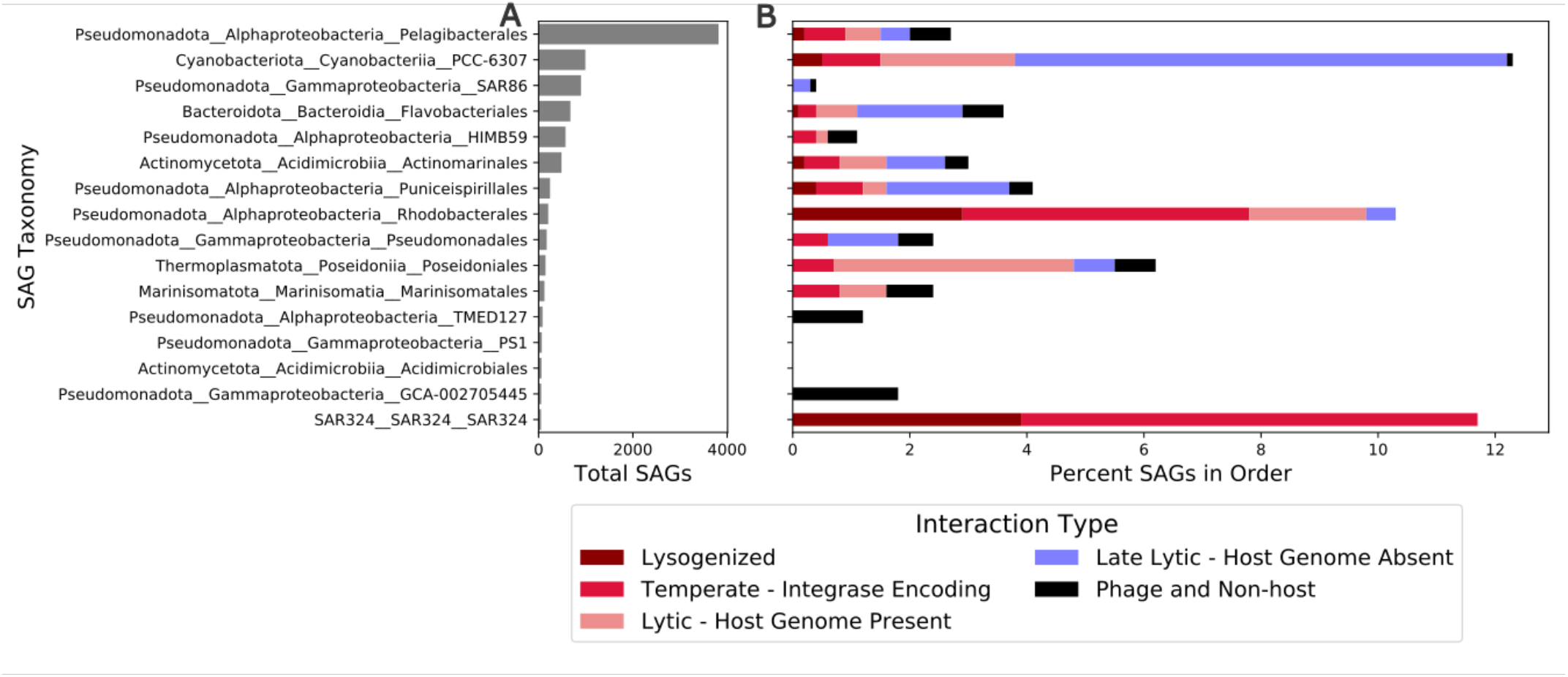
Abundance and characteristics of infection across taxa with at least 50 representatives in GORG-Tropics. A. Number of SAGs elonging to order-level taxa displayed as phylum-class-order. B. Percent cells containing a phage with colors indicating the type of phage nteraction observed.

Pelagibacterales exhibited significantly lower than expected frequencies of phage interactions (2%, Table 1, Figure S3A). Other abundant, slow-growing groups including SAR86 and HIMB59 exhibited even lower frequencies of phage interactions despite being abundant in the GORG-Tropics database (0.3% and 0.5% of recovered cells per order). These groups represent the most common microbial taxa throughout the global surface ocean [16,57], and these taxa are known to be slow growing with low to undetectable rates of respiration [51] and streamlined genomes [58,59]. These results suggest that low rates of phage infection may contribute to the numerical abundance of these prokaryoplankton groups.

### Phage-host interaction characteristics across host populations

To better understand the nature of phage infections among microbial populations, we examined infection strategies of phages detected in all prokaryotic SAGs. Specifically, we compared phage co-assembly with host (aka lysogeny) versus all other interactions, and phage-only SAGs (cells experiencing late-lytic infection) versus co-occurrence of phage and host genomic sequences within SAGs (earlier stages of lytic infection or lysogeny), both across order-level taxonomic groups with at least 10 phage interactions identified. This revealed significant differences in the distribution of infection characteristics (Table S1, Figure 2B, Figures S2B and S2C) and a decoupling between infection strategy and infection frequency.

Alphaproteobacterial orders represented some of the most infected (Rhodobacterales) and least infected (Pelagibacterales, HIMB59) prokaryoplankton taxa (Figure 2B). Yet, across these groups, host genomic sequences were more frequently found in SAGs that also contained phage sequences, and in the case of Rhodobacterales, their phage sequences were more frequently integrated with the host’s genome (Figure 2B, S2C). This suggests that observed phage infections of Alphaproteobacterial SAGs are less lytic and more likely to be temperate, or chronic. By contrast, phages of *Prochlorococcus* (Cyanobacteria), and Flavobacterales (Bacteroidia) -- orders that also differ in phage interaction frequencies -- were more frequently found in SAGs without host genomic sequences, indicating many infected SAGs were in the late stages of lytic infection cycles (Figure 2B, S3C). Phages predicted to infect these hosts were also less likely to be integrated with host genomic sequence (Figure S3B), together suggesting that phages of these hosts more frequently utilize a lytic viral infection strategy.

### Phage associations within redox-active prokaryoplankton

For the most deeply sampled station, SWC-09, SAGs were created from both redox-active and randomly sampled prokaryoplankton particles, enabling comparison of phage interactions between these two populations. A similar proportion of randomly sampled SAGs (3.3%) and redox-active SAGs (3.2%) were found to be associated with a phage at SWC-09, indicating that at the community level, similar numbers of redox-active and total cells contained phage genomic material, similar to prior observations [51]. The distribution of infection states differed between populations, with a higher proportion of interacting phages either lysogenized, encoding an integrase, and co-occurring with host genomic DNA in the redox-active cells, and greater proportion of lytic interactions within randomly sampled cells, potentially reflecting lower redox-activity within cells undergoing later stages of lytic infection and/or enrichment of lysogeny within redox-active cells (Figure S4). Interestingly, a small number of redox-active cells with only phage genomic material present were observed. For example, 5 redox-active cells containing phage sequences and no host genomic information were predicted to infect the order Punicispirialles (Figure S4B). This suggests that redox activity is sustained during the later stages of phage infection in some cases.

### Phages found within non-host cells

Computational host prediction methods identified putative hosts for 474 out of the combined 545 identified phages identified within redox-active and randomly sampled SAGs, with the most common predicted hosts being *Prochlorococcus* (160 phages), *Pelagibacter* (102 phages), Flavobacteriaceae (44 phages), Rhodobacteriaceae (27 phages), Puniceispirillales (25 phages), and Actinomarinales (12 phages). There were 63 cases in which the predicted phage host did not match the taxonomy of the cell the phage was found within (Table 1, Figure S3A). Flavobacteriales phages (16) and cyanophages (18) were most commonly found within non-host cells, with cyanophages found in the greatest diversity of cells (Figure 3A). Some of these mismatches between the predicted and the observed phage hosts may be due to errors in computational host prediction, e.g., due to insufficient coverage of marine viral diversity by genome reference databases. However, many of the computationally predicted cyanophages found in SAGs of non-photosynthetic bacterial lineages encoded photosynthetic auxiliary metabolic genes, which provides additional support to the computational host prediction (Figure S5).

**Figure 3.**
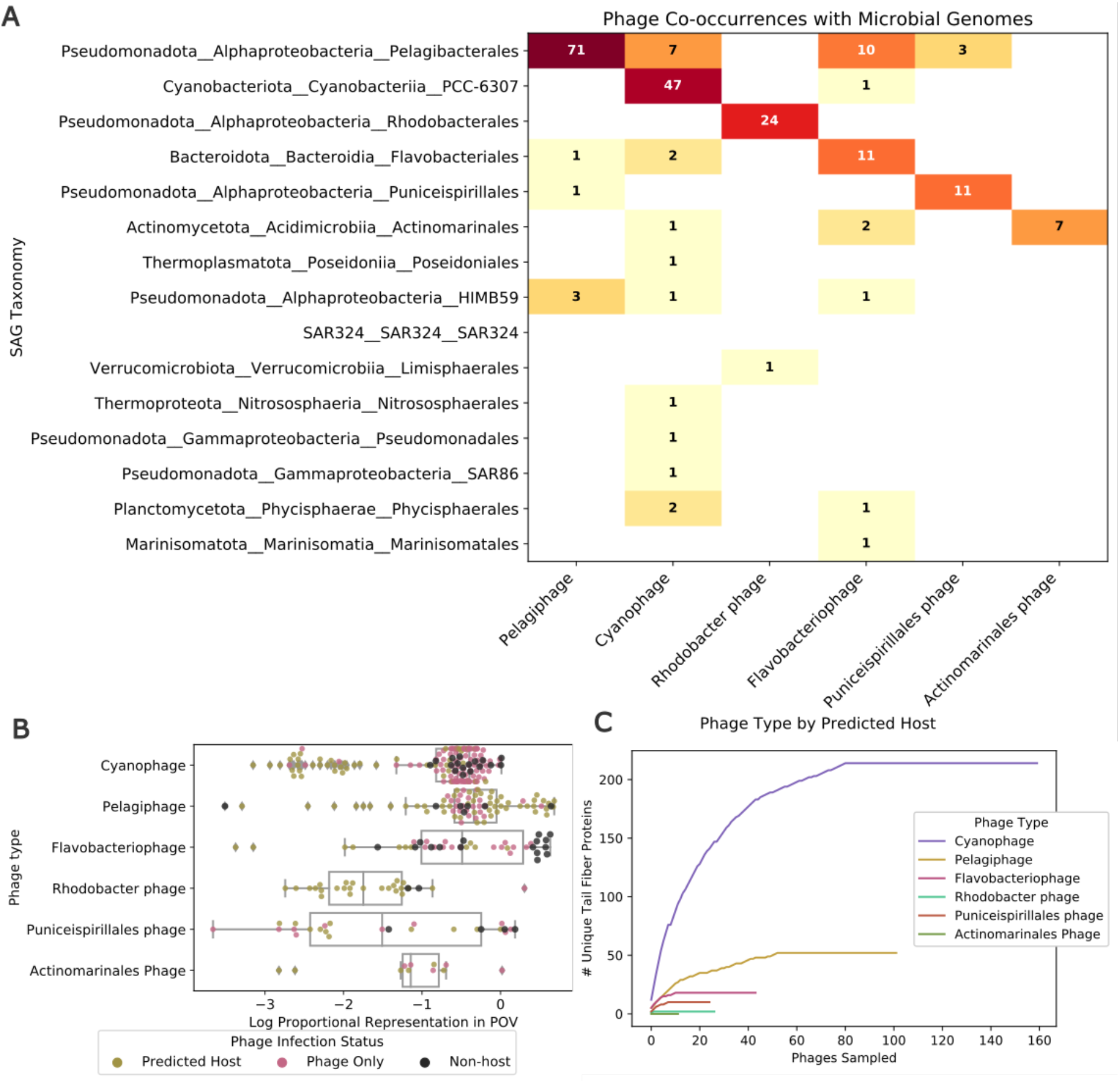
Distribution of phages across SAGs. A. Plot depicting distribution of most common phage groups across most common order-level taxa within GORG-Tropics (taxa with 50 or greater representatives). B. Log relative recruitment of metagenomic reads from the Pacific Ocean Virome (POV) to GORG-Tropics phages, separated by predicted host with overlaid points indicating infection status of cell from which phage was recovered. C. Collectors curve of unique tail fiber proteins sampled from phages with the most to least number of tail fiber proteins per phage for phages of the top five most predicted phage hosts.

Some of the observed non-host interactions may be due to accidental co-sorting of cells and free viral particles during SAG generation. Yet, based on FACS droplet sizes and viral and bacterial abundances in marine surface waters, the predicted frequency of accidental co-sorting under the applied conditions was less than 1 in 2,500 [13]. This translates to about five expected accidental free particle co-sorts in the entire GORG-Tropics collection, which is substantially less than the observed 63 cases of virus-host mismatches. In addition, if these instances were due to random events during sorting, we would expect to see a distribution of non-host viruses in SAGs that mirrors their particle abundances in nature. To test this, we compared the relative recruitment of reads from the Pacific Ocean Virome (POV) to phage genomes from GORG-Tropics. Phages of *Pelagibacter* exhibited significantly higher abundances than those of *Prochlorococcus* (p= 5.97e-05, Figure 3B) despite phages of *Prochlorococcus* being more frequently observed in non-host SAGs, indicating that extracellular abundance is likely not an indicator of the propensity to engage in non-host interactions. The distribution of non-host phages across microbial groups was also non-random, with cyanobacterial SAGs having a significantly lower frequency of such associations in comparison to other lineages (Figure 2B, Table S1). Given this evidence, we conclude that co-sorting is likely not the primary explanation for the discrepancies between the predicted and observed phage-host interactions.

Some cyanophages are known to have broad host ranges that span multiple cyanobacterial species [60,61] and even both marine and freshwater hosts [62], while other cyanophages have narrow host ranges [63]. Many cyanophages recovered from GORG-Tropics were enriched in genes encoding receptor-binding proteins (Figure 3C), suggesting permissive attachment strategies that could lead to non-specific attachment. Obligately lytic phages depend on cellular entry to propagate, so in environments where host populations are especially microdiverse and/or sparse, permissive attachment may increase the probability of finding a host.

The distribution of apparently non-infecting phages within SAGs of heterotrophic taxa and the near-absence of non-infecting phages within phototrophs (*Prochlorococcus*) suggests that the presence of non-infective phage-microbe interactions may be related to host trophic status. Attracting non-infecting viruses could be a strategy used by heterotrophs for nutrient acquisition in oligotrophic waters, as has been previously hypothesized through modelling [64] and observed using laboratory experiments [65]. Furthermore, these results indicate that phages of different host groups may differ in their propensity to engage in non-host entry.

The occurrence of phages expected to infect one host but found in a different cell type has been previously observed within a smaller collection of marine bacterial SAGs: two phages with strong homology to *Synechococcus* phages were found in SAGs of the marine *Roseobacter* clade. Subsequent experiments challenging a cultured *Roseobacter* with a *Synechococcus* phage did not cause *Roseobacter* to lyse [13], suggesting that observed interactions are non-lethal. A recent preprint also identifies an estuarine cyanophage with a broad interaction range beyond its known cyanobacterial host [66].

If a non-specific phage enters a cell while it is experiencing infection by a virulent phage, entry of non-killing phages would provide a mechanism for gene exchange among phages with non-overlapping host ranges [67,68]. Cryptic entry, combined with recombination with virulent phages also provides a mechanism for gene exchange across phages targeting distant hosts [67] as well as “host swapping”, hypothesized to occur based on gene content similarity between phage isolates capable of infecting phylogenetically distinct hosts [69].

### Relatedness of identified phage genomes and potential for genomic exchange

To look more closely into the genomic relatedness of phages found in GORG-Tropics SAGs and in the context of previously characterized phage genomes, their protein sequences were clustered with phage genomes extracted from GenBank by INPHARED ((Cook et al. 2021), downloaded May 11, 2023) using vContact2 (Bolduc et al. 2017). This resulted in a large network of phage genomes, with each genome connected to another based on shared protein clusters. Of the GORG-Tropics phages, 216 (38%)) fell into 57 genus-level virus operational taxonomic units (vOTUs), 50 (9%) were singletons that are excluded from the vContact network. Others were included in the vContact network but were not assigned a vOTU (195; 34%) or spanned more than vOTU (37 ;6.0%). Despite the large number of viruses falling outside of vContact2-defined genus-level vOTUs, the majority of verified phages from GORG-Tropics included in the network were located within one region, connected to each other and other marine phages (Figure 4A). We next examined a subnetwork of GORG-Tropics phages, their immediate neighbors, and all connections therein, to see how GORG-Tropics contributes to network structure (Figure 4B). This subnetwork includes INPHARED-sourced marine phage genomes such as T4-like phages infecting picocyanobacteria (*Kyanoviridae)*, phages of Pelagibacterales (*Pelagibacter phage Mosig* [70], *Pelagibacter phage HTVC008M* [71]), and phages of gammaproteobacterial clade OM43 (*Methylophage Melnitz* [69]). This subnetwork also includes marine viruses belonging to the *Autographiviridae*, including cyanophages and other marine T7-like phages such as *Pelagibacter phage HTVC010P*. Without GORG-Tropics viruses, T4-like marine phages, including the *Kyanoviridae*, formed a tight cluster within the subnetwork separate from the *Autographiviridae* (Figure 4C). The inclusion of GORG-Tropics phages connected marine T4-like phages, including *Kyanoviridae*, to other members of the subnetwork (Figure 4B), making the overall subnetwork more connected, as evidenced by a decreased network radius from 15 to 12. GORG-Tropics viruses increased network heterogeneity and decreased the number of total network components, all showing that viruses from GORG-Tropics connect what would otherwise be interpreted as discrete phage groups. This result highlights that gene exchange within wild phage populations extends beyond the taxonomic boundaries defined largely by phage genomes sourced from isolated, lytic phage strains, and that as more phages are sampled, taxonomic boundaries will continue to shift.

**Figure 4.**
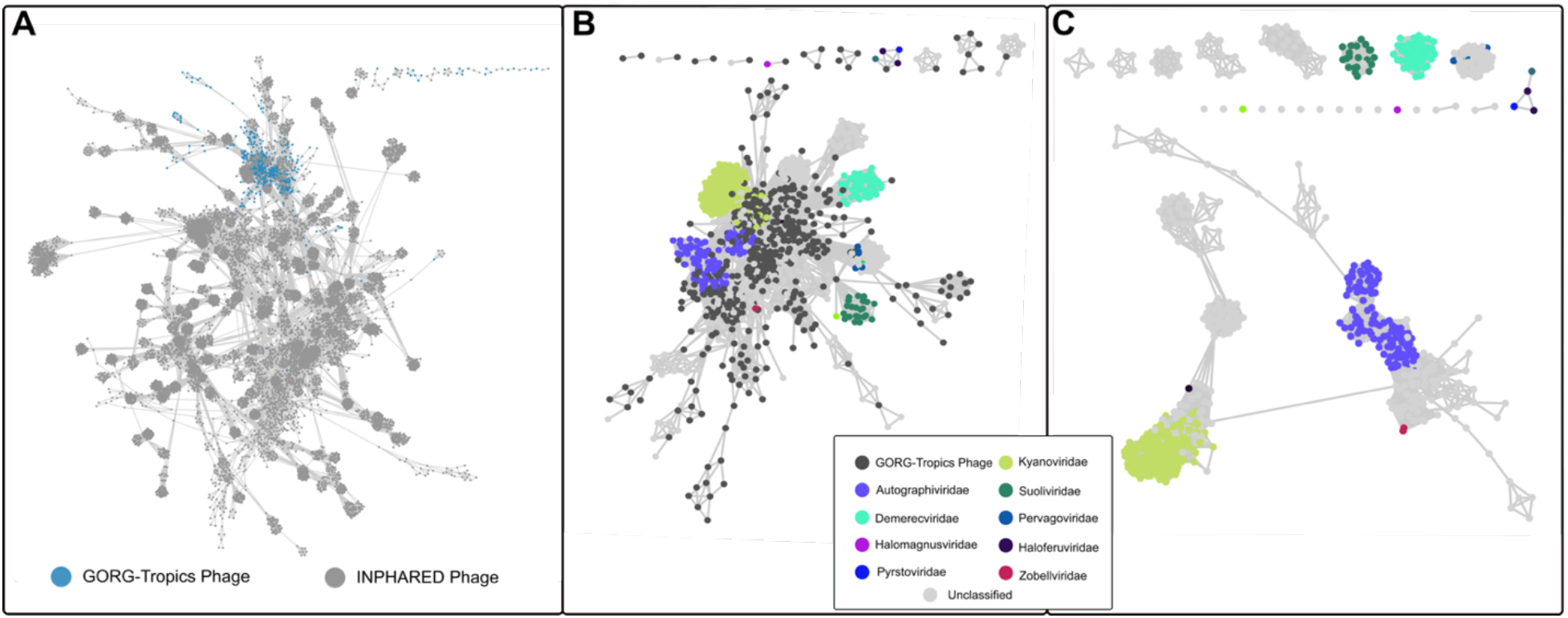
Phage gene sharing networks. vContact network showing Genbank phages collected by INPHARED and GORG-Tropics viruses. A. Nodes represent viral genomes, edges represent vContact-identified shared protein content between viruses. Circles are viral nodes collected by INPHARED, triangles are viral genomes extracted from GORG-Tropics SAGs. B. Subnetwork of nodes representing GORG-Tropics viruses and immediate neighbors. Nodes are colored by currently available information about viral family membership. C. Subnetwork of immediate neighbors of GORG-Tropics viruses redrawn with GORG-Tropics viruses removed.

The increased network connectivity facilitated by GORG-Tropics phages is driven by low edge-weight connections, indicating that the connected phages contain a small number of shared genes. While the majority of genes shared between phages with low edge weights had <50% amino acid identity, 3.1% of weakly connected phage genomes from GORG-Tropics (81 out of 2,588 network edges with an edge weight <10) shared at least one protein with ≥95% amino acid identity. Of those cases, 10 involved phage pairs predicted to infect hosts of divergent taxonomic Orders. Furthermore, we found entire protein-encoding gene cassettes containing shared genes with varying amino acid identities between phages predicted to infect divergent hosts. For example, a predicted *Prochlorococcus* phage, phage_AG-359-J16, containing 44.6% of annotated genes with closest matches to cultivated cyanophages, had a RecA- encoding contig with nine protein-encoding genes syntenic to *Pelagibacter* phages phage_AG-920-F02 and phage_AH-324-B02 with amino acid identities ranging from 35 to 96% (Figure 5). These examples illustrate cases of genomic exchange, which is known to be common in many viral groups [67,72]. Because free phage particles are encapsulated and their genomes protected, the only opportunity for genetic exchange is within cells. Therefore, the observed co-occurrence of multiple phages in individual prokaryoplankton cells represented by GORG-Tropics SAGs, as well as phages identified within non-hosts, reveals an important mechanism for genomic exchange among phages that infect divergent hosts.

**Figure 5.**
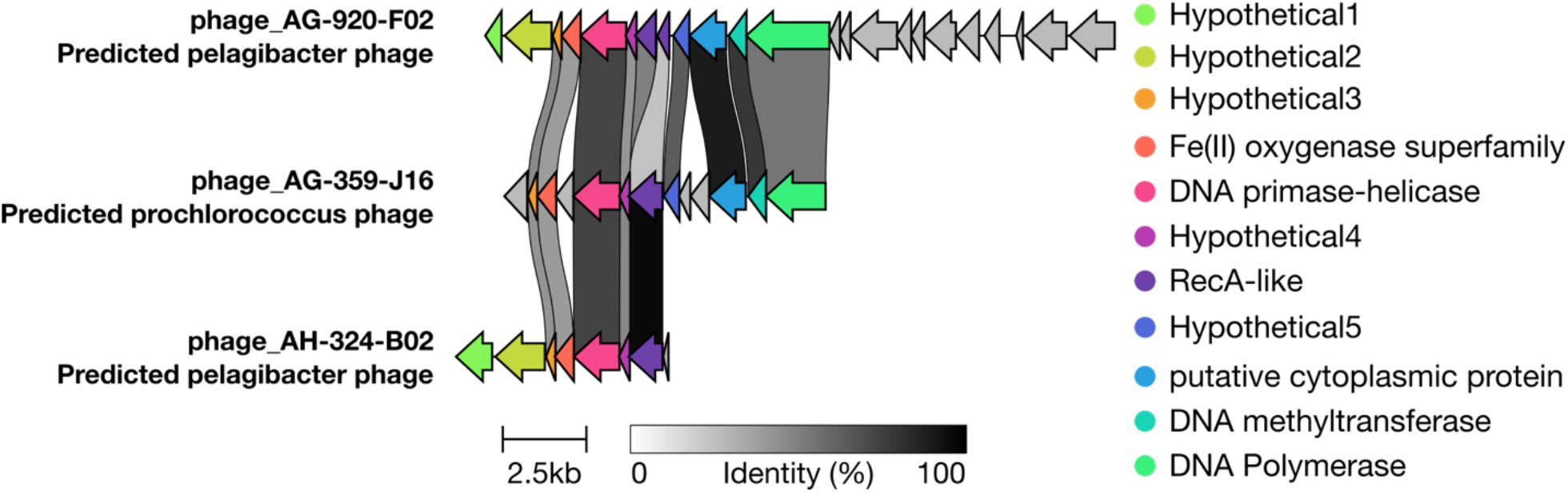
Gene sharing between GORG-Tropics phages. Alignment of contigs from two Pelagibacter phages and a Prochlorococcus phage recovered from GORG-Tropics SAGs. Arrows represent coding sequences on genomic contigs from each predicted phage, arrows of the same color represent homologous genes, connecting grey regions represent shared amino acid identity between aligned genes.

## Conclusions

By examining the composition of viral sequences in GORG-Tropics, a global collection of 12,715 tropical and subtropical epipelagic marine prokaryoplankton SAGs, we obtained one of the first quantitative assessments of viral interactions with marine prokaryoplankton at single-cell resolution. Our estimate of 4.2% GORG-Tropics SAGs containing a phage (ranging from 1 to 19% among sampled sites) is in agreement with prior studies in the oligotrophic marine epipelagic zone using different methods, and the highest rates observed here were in samples from high productivity regions. However, these rates may be underestimates due to the conservative approach we took in the separation of phage sequences from other mobile genetic elements and the potential presence of novel viruses that could not be recognized and validated with current bioinformatics tools. An additional source of potential underestimates includes the removal of < 10 kbp viral genomes in our computational workflow to minimize false positives [73] but this may have precluded the detection of ssDNA and other small viruses. Furthermore, the DNA amplification and sequencing techniques that were used to generate GORG-Tropics SAGs were not designed to recover sequences of RNA viruses and, potentially, some DNA viruses containing nucleotide modifications or strong bonds with proteins and other molecules. Thus, the observed frequency of prokaryoplankton-virus interactions should be viewed as dsDNA phage-specific and conservative.

The highest and the lowest frequencies of viral associations were observed in abundant prokaryoplankton lineages previously observed to be the most metabolically active (Cyanobacteria and Rhodobacterales) and the least metabolically active (Pelagibacterales and SAR86), respectively. Meanwhile, infection strategy (host lysogeny versus lytic infection) varied by host’s taxonomy regardless of host’s abundance and activity, with viruses of abundant Alphaproteobacteria lineages Rhodobacterales, Pelagibacterales, HIMB59, and Puniceispirillales more frequently integrated into the host’s genome and more frequently containing genomic indicators of lysogeny than viruses of *Prochlorococcus* (Cyanobacteria) and Flavobacteriales (Bacteroidia). Furthermore, we identified many instances in which the observed virus-cell associations differed from the virus’s computationally predicted host, suggesting that a subset of marine phages enter non-host cells. The observed cases of co-occurrence of both putatively virulent and non-infective viruses in the same cell and the finding of highly similar genome regions in phages with divergent hosts suggest that non-infective phage entries contribute to exchange of genomic information across diverse phages. Collectively, this study demonstrates a new approach to obtain large-scale, quantitative data on viral interactions with cellular microorganisms in nature, contributing new knowledge in microbial ecology and evolution.

## Acknowledgements

We thank Maria Pachiadaki (Woods Hole Oceanographic Institute) and Jacob Munson-McGee for insightful advice and comments. This work was supported by the Simons Foundation (Life Sciences Project Award ID 510023 and 1829879 to R.S.).

---

**Figure S1.**
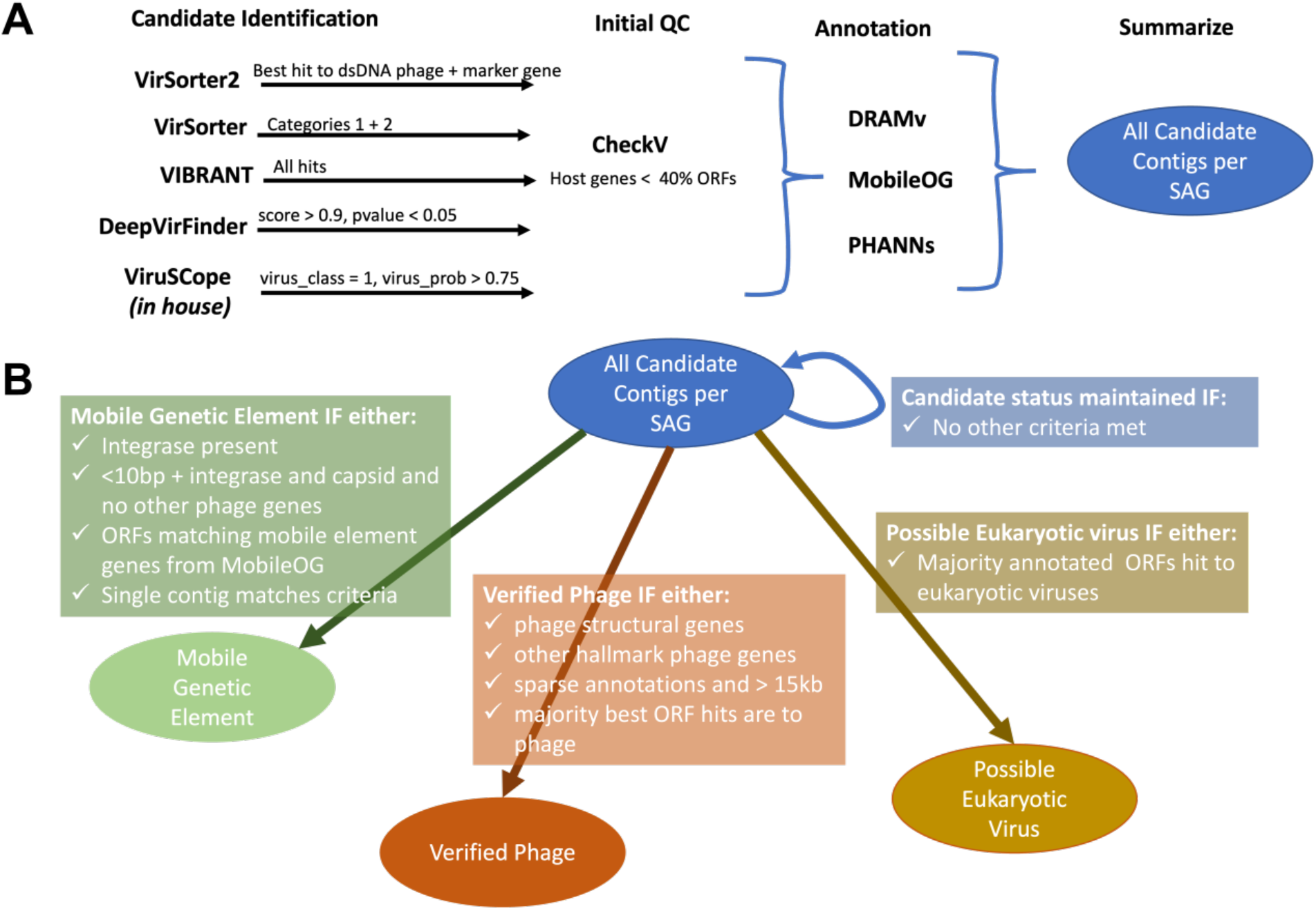
Caption. Phage Identification and Confirmation. Workflow for initial identification of virus candidate contigs (A). Considerations for sorting all viral candidate contiguous sequences recovered from SAGs into categories. Only results for sequences falling into the ‘Verified Phage’ category were included in subsequent analyses presented in this manuscript (B).

**Figure S2.**
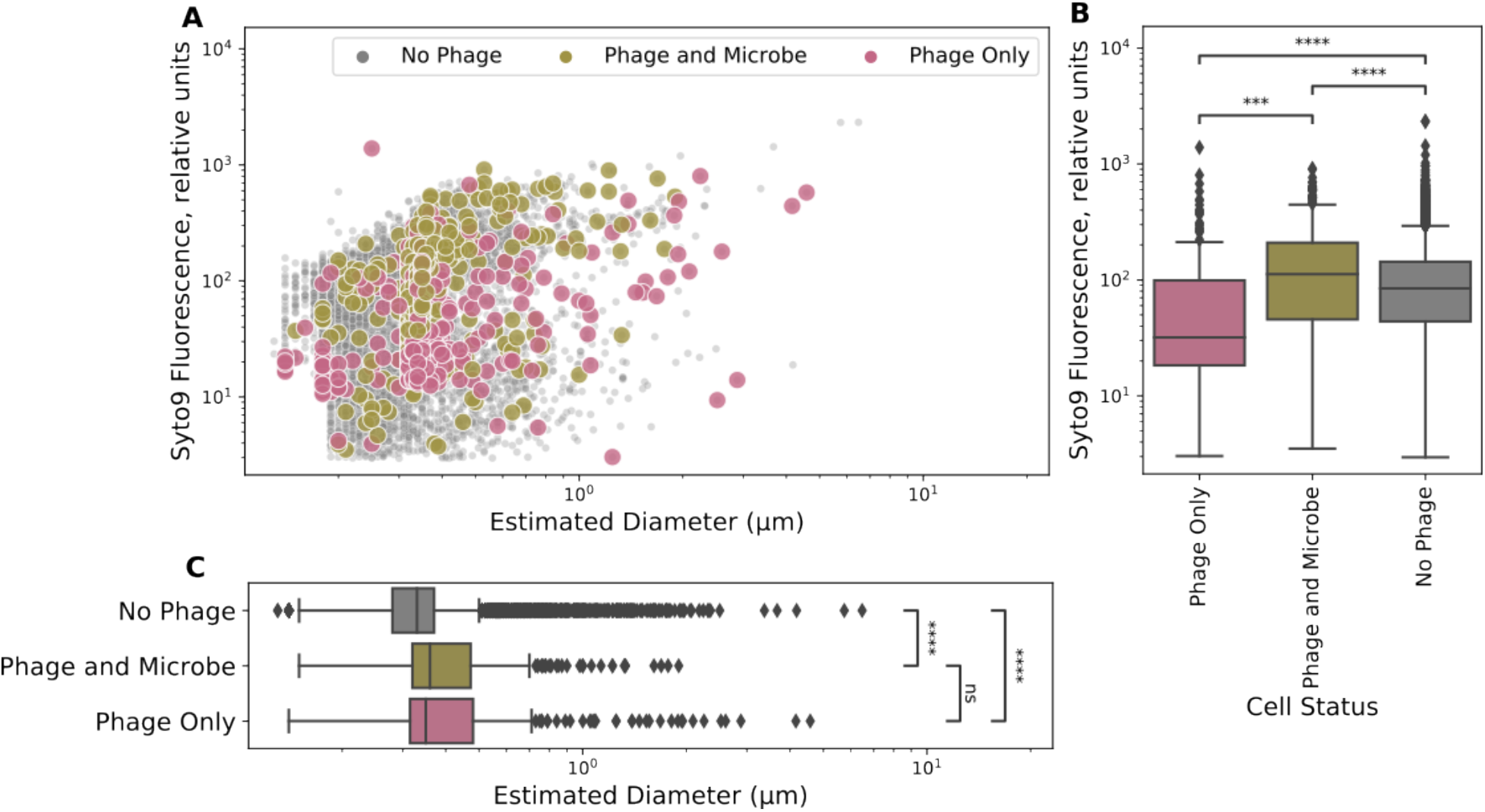
Caption. Phenotypic properties of infected and uninfected cells. A. Comparison of Syto9 fluorescence against estimated cell diameter for the GORG-Tropics collection of cells. Each point represents a sorted particle associated with a SAG. Particles are colored based on virus status of the SAG. B. Comparison of Syto9 Fluorescence for SAGs of differing infection status. C. Comparison of estimated cell diameter for cells of differing infection status. Significance brackets in plots B and C indicate that a Kruskal-Wallis test identified a significant difference between groups, p << 0.0001, brackets specifically indicate results of Dunn’s post hoc tests with bonferroni correction. P-values are noted as follows: ns: not significant; *: p <= 5.00e- 02; **: p <= 1.00e-02; ***: p <= 1.00e-03; ****: p <= 1.00e-04.

**Figure S3.**
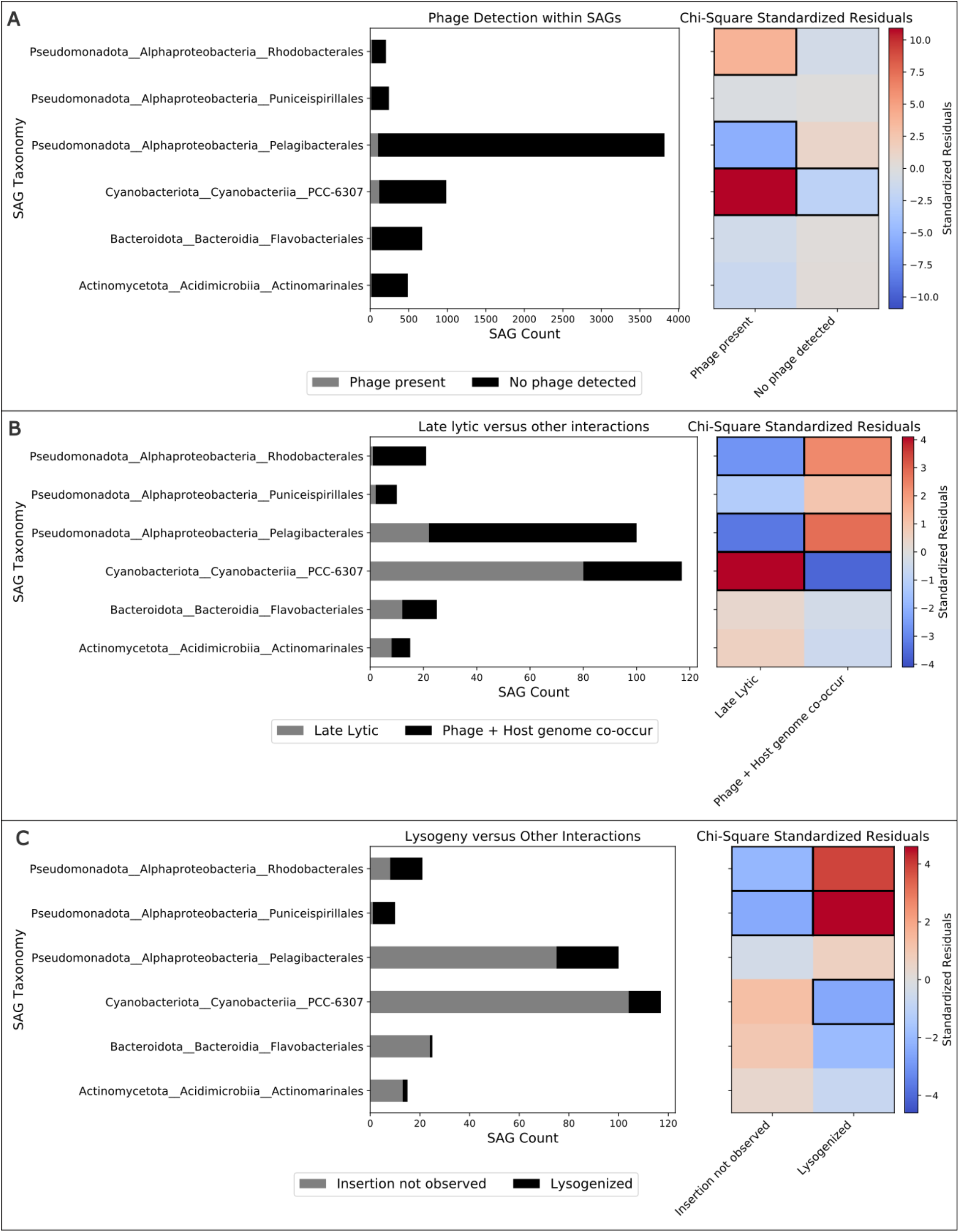
Caption. Infection characteristics for order-level hosts. Orders with at least 10 phage interactions and non-zero values for the categories tested are included in each plot. A. Plot of the prevalence of phage interactions for order-level taxa with at least 10 observed interactions (left) with standardized residuals of Chi2 test of independence of infection by SAG taxonomy (right). B. Plot of distribution of Late-lytic versus host-genome-associated phage-microbe interactions within the same six taxonomic groups, displaying the proportion of phage interactions in which only viral sequence material was identified (Late Lytic) and interactions where both phage and microbial genomic sequence information was observed (Phage and Host genome co-occur) and associated standardized residuals from a Chi2 test of independence. C. Plot of distribution of interactions where lysogeny was observed (Lysogenized) versus phage-microbe interactions in which it was not (Insertion not observed) and associated standardized residuals from a Chi2 test of independence. For standardized residuals plot, standardized residual values indicating significantly different distributions from expected are highlighted with a solid square.

**Figure S4.**
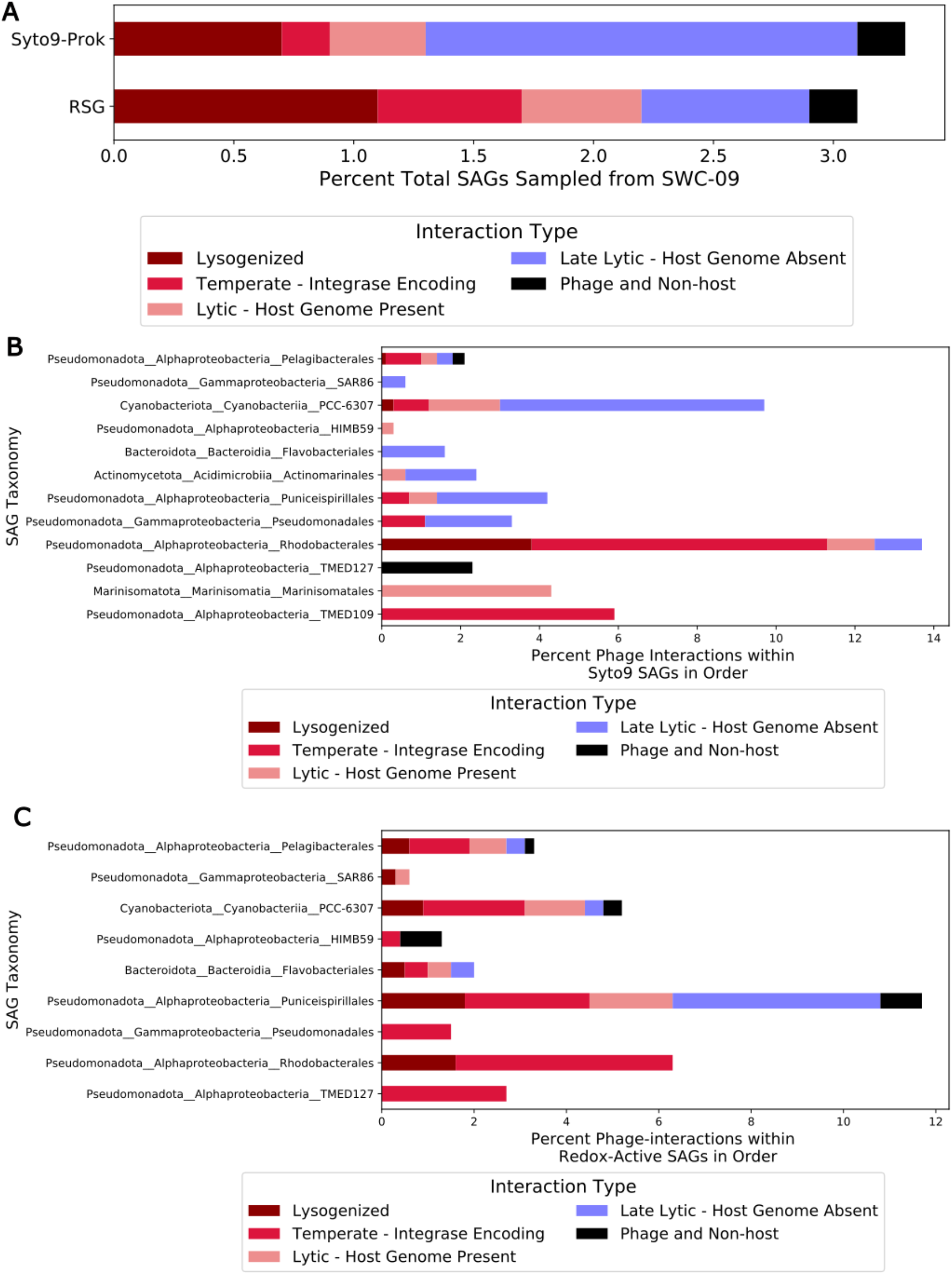
Caption. Distribution of infection strategies in randomly sampled and redox-active prokaryotic populations from sample SWC-09. A. Overall distributions of infection strategies from phage-containing SAGs. Percent cells containing a phage with colors indicating the type of phage interaction observed for B. randomly sampled SAGs (selected based on Syto9 fluorescence) and C. redox-active SAGs selected based on RSG fluorescence.

**Figure S5.**
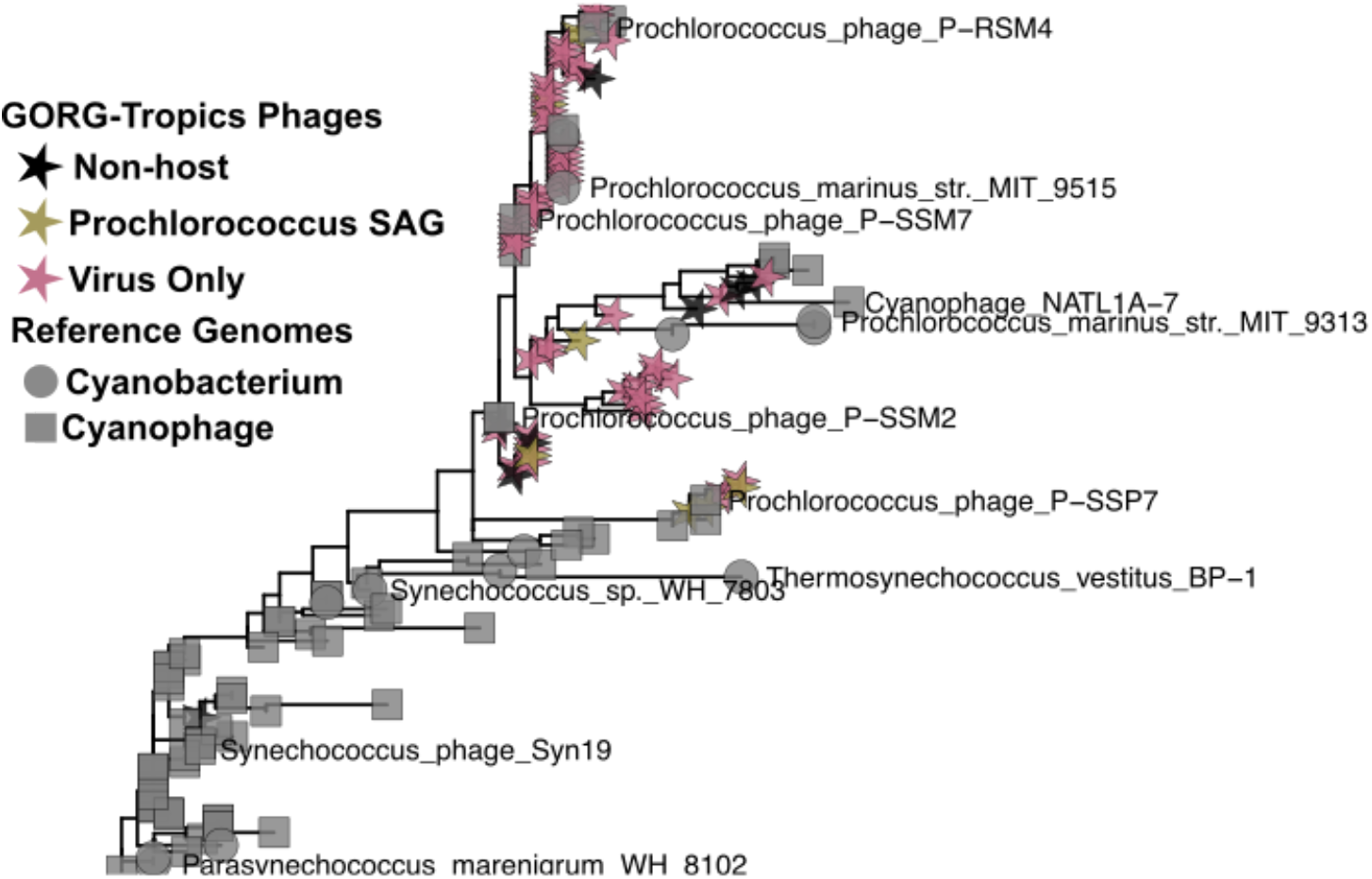
Phylogenetic relationship of PsbA genes. recovered from phages from GORG-Tropics SAGs with PsbA from marine Prochlorococcus and Synechococcus, and other cyanobacterial phages.

**Table S1.** Chi2 test statistics for tests of independence of predicted phage host by categorical variables related to infection strategy for the six host groups for which at least 10 phages were recovered from randomly samples SAGs prokaryotic SAGs within GORG-Tropics. For comparison of distribution of non-host phages, only SAGs for which host genomic information was available was included, reducing the number of taxonomic groups with at least 10 phages to 5, explaining the lower DoF for this Chi2 test.

| Variable | DoF | N | Chi2 Statistic | p-value |
| --- | --- | --- | --- | --- |
| Contains Phage | 5 | 6414 | 175.31 | 5.34E-36 |
| Infection Strategy -<br>Lysogenized phage | 5 | 288 | 60.66 | 8.86E-12 |
| Infection Strategy – Late-lytic phage | 5 | 288 | 64.163 | 1.67E-12 |
| Microbe Contains Non-host<br>Phage | 4 | 164 | 21.58 | 0.0002 |

